# Electron-counting MicroED data with the K2 and K3 direct electron detectors

**DOI:** 10.1101/2022.07.04.498775

**Authors:** Max T.B. Clabbers, Michael W. Martynowycz, Johan Hattne, Brent L. Nannenga, Tamir Gonen

**Affiliations:** Department of Biological Chemistry, University of California, Los Angeles CA 90095; Howard Hughes Medical Institute, University of California, Los Angeles CA 90095; Chemical Engineering, School for Engineering of Matter, Arizona State University, Tempe, AZ; Center for Applied Structural Discovery, Biodesign Institute, Arizona State University, Tempe, AZ; Department of Physiology, University of California, Los Angeles CA 90095

## Abstract

Microcrystal electron diffraction (MicroED) uses electron cryo-microscopy (cryo-EM) to collect diffraction data from small crystals during continuous rotation of the sample. As a result of advances in hardware as well as methods development, the data quality has continuously improved over the past decade, to the point where even macromolecular structures can be determined *ab initio*. Detectors suitable for electron diffraction should ideally have fast readout to record data in movie mode, and high sensitivity at low exposure rates to accurately report the intensities. Direct electron detectors are commonly used in cryo-EM imaging for their sensitivity and speed, but despite their availability are generally not used in diffraction. Primary concerns with diffraction experiments are the dynamic range and coincidence loss, which will corrupt the measurement if the flux exceeds the count rate of the detector. Here, we describe instrument setup and low-exposure MicroED data collection in electron-counting mode using K2 and K3 direct electron detectors and show that the integrated intensities can be effectively used to solve structures of two macromolecules between 1.2 Å and 2.8 Å. Even though a beam stop was not used in these studies we did not observe damage to the camera. As these cameras are already available in many cryo-EM facilities, this provides opportunities for users who do not have access to dedicated facilities for MicroED.

## 1. Introduction

Microcrystal electron diffraction (MicroED) uses continuous rotation data collection for protein structure determination from three-dimensional (3D) crystals (Nannenga *et al*., 2014). The sample preparation and diffraction experiment are similar to electron cryo-microscopy (cryo-EM), whereas the data collection strategy, data processing, and structure refinement are analogous to macromolecular X-ray crystallography. Electrons interact strongly with the electrostatic potential of the crystal and have a relatively high ratio of elastic to inelastic interactions (Henderson, 1995). Therefore, MicroED can be used to determine structures from very small crystals even at very low exposures. The signal in a diffraction pattern can be improved by collecting data from a larger sample with more scattering unit cells. Because electrons interact strongly with matter, larger crystals introduce problems as the probability for multiple interactions and absorption increases (Cowley & Moodie, 1957). Alternatively, the signal can be boosted by increasing exposure rate or time, but this will also increase radiation induced damage to the sample, limiting the useful structural information that can be obtained from crystalline biological specimen (Hattne *et al*., 2018). Crystal thickness should therefore be optimized for obtaining the highest signal to noise with a minimal exposure (Martynowycz *et al*., 2021), and for this a sensitive and fast camera is key. For a given crystal size, a sensitive camera allows the exposure to be reduced, thus limiting radiation damage without sacrificing the integrity of the measurement.

Scintillator based complementary metal-oxide semiconductor (CMOS) cameras offer reasonable frame rates and sufficient signal-to-noise ratio when shutterless mode is used for continuous rotation MicroED (Nannenga *et al*., 2014). These cameras record data using integrating mode, where the number of electrons is determined by the charge accumulated in a pixel during a readout-cycle. Whereas charge-coupled device (CCD) cameras also have been effectively used for protein structure determination of similar sized crystals, overall they tend to be less sensitive, slower and more involved to operate (Yonekura *et al*., 2015; Zhou *et al*., 2019). Hybrid pixel detectors have a high dynamic range for diffraction experiments, but have relatively large pixels and small arrays that may make them unsuitable for large unit cells (Nederlof *et al*., 2013; Clabbers *et al*., 2017).

In single particle cryo-EM imaging, direct electron detectors are commonly used as these have a high detective quantum efficiency and small pixels combined with a large effective field of view (Li, Mooney *et al*., 2013; Li, Zheng *et al*., 2013; Nakane *et al*., 2020). However, their use in diffraction has been limited, mainly due to concerns regarding the frame rate and the limits it imposes on the dynamic range. The dynamic range of a counting detector is determined by the frame rate, as the number of electrons that can be accurately measured on a single frame is limited, such that often only the first electron that arrives on a pixel during a readout-cycle is reported. Any subsequent electrons on the same pixel during the same readout-cycle are ignored, leading to coincidence loss. These pileup effects mostly affect strong, low-resolution reflections where they can severely corrupt the measurement of their intensities. Regardless of these drawbacks, sensitive detectors using electron counting are highly desirable for MicroED as they can help to accurately record even the very weak intensities using a highly attenuated exposure.

For these reasons, previous MicroED studies using direct electron detectors such as the Falcon 3 (Thermo Fisher) and the DE64 (Direct Electron) for protein structure determination relied on integrating/linear mode rather than counting (Hattne *et al*., 2019; Takaba *et al*., 2021). It is, however, possible to collect electron-counted MicroED data by significantly lowering the exposure, and this was recently shown to provide data of sufficiently high quality for *ab initio* macromolecular structure determination using the Falcon 4 (Thermo Fisher) direct electron detector (Martynowycz *et al*., 2022; Clabbers *et al*., 2022). These results are encouraging, however MicroED data collection using direct electron counting is not yet mainstream despite the availability of these cameras in cryo-EM laboratories. The most common direct electron cameras at cryo-EM centers are still the K2/K3 Gatan direct electron detectors and thus far these have been recalcitrant to MicroED in counting mode.

Here, we determine macromolecular structures using continuous rotation MicroED data collection on K2 and K3 direct electron detectors (Gatan) operated in electron counting mode. We describe the adjustments in sample preparation and hardware setup that were necessary to accommodate low exposure data collection in counting mode and characterize the performance of these cameras. These results suggest that electron-counting can be routinely used for protein structure determination by MicroED, making any diffraction applications more accessible to a wider audience as these cameras are generally available to structural biologists using shared cryo-EM facilities.

## 2. Materials and Methods

### 2.1 Crystallization

#### 2.1.1 Crystallization of proteinase K

For experiments on the K2, crystals of proteinase K were grown by hanging drop by adding 2 μl of precipitant solution containing 1.2 M ammonium sulfate, 0.1 M Tris pH 8.0 to 2 μl of 50 mg/ml of proteinase K from *Engyodontium album*. The drops were equilibrated over 500 μl of precipitant in the wells and incubated at room temperature overnight. Crystals of proteinase K for data collection on the K3 were grown by dissolving 40 mg/ml protein in 20 mM MES-NaOH pH 6.5. The protein solution was mixed at a 1:1 ratio with a precipitant solution of 0.5 M NaNO_3_, 0.1 M CaCl_2_, 0.1 M MES-NaOH pH 6.5. The mixture was incubated at 4 °C and crystals grew overnight.

#### 2.1.2 Crystallization of triclinic lysozyme

Crystals of hen egg-white lysozyme (*Gallus gallus*) were grown by dissolving 10 mg/ml protein in a filtered solution of 0.2 M NaNO_3_, 0.05 M NaAc pH 4.5, The mixture was incubated overnight at 4 °C and crystals grew after further incubation at room temperature for one week.

### 2.2 Grid preparation

Standard holey carbon grids (Quantifoil, Cu 200 mesh, R2/2) were glow discharged for 30 s at 15 mA on the negative setting. For each sample, 3 μl of crystalline slurry was deposited onto a grid, excess liquid blotted away from the back side, and the sample was then rapidly vitrified in liquid ethane using a Leica GP2 plunger freezer set at 4 °C and 90% humidity. Grids were stored in a liquid nitrogen dewar prior to use.

### 2.3 Focused ion beam milling

Crystals of lysozyme and proteinase K were thinned prior to MicroED experiments using the K3 camera. Grids were loaded into an Aquilos (Thermo Fisher) focused ion-beam and scanning electron microscope (FIB/SEM) and crystalline lamellae were prepared as described previously (Martynowycz *et al*., 2021). Briefly, grids were sputter coated to cover the sample in an approximately 100 nm thick layer of platinum. Crystals were first thinned using a coarse milling procedure to a thickness of 2 μm at a current of 0.3 nA. Afterwards, cleaning cross sections at a lower current of 50 pA were used to thin the lamellae to about 300 nm thickness. Grids were transferred directly after milling to the transmission electron microscope (TEM) and rotated by 90° relative to the milling direction such that the rotation axis of the TEM is perpendicular to the milling direction.

### 2.4 Data collection

#### 2.4.1 MicroED data collection using electron counting on the K2

MicroED data were collected on a Titan Krios TEM (Thermo Fisher) operated at 300 kV. The K2 (Gatan) director electron detector (5 μm pixel size, 3,840×3,712 pixels) is retractable and positioned on-axis at the bottom of the column. The camera operates at an internal frame rate of 400 Hz, and minimal coincidence loss below an exposure of about 8 e^-^/pixel/s (Li, Mooney *et al*., 2013). For low exposure data collection, the microscope was set up using a 50 μm C2 aperture and a spot size of 11. The beam was spread under parallel conditions to 15 μm diameter, corresponding to an exposure rate of about 0.01 e^-^/Å^2^/s. A 200 μm selected area (SA) aperture was used to select diffraction from a 4 μm diameter area on the crystal. Diffraction data were recorded using 2× binning at a readout rate of 40 frames per second. Data were collected with a rotation speed of 1.52° per second.

#### 2.4.2 MicroED data collection using electron counting on the K3

MicroED data were collected on a Titan Krios TEM (Thermo Fisher) operated at 300 kV and equipped with a K3 direct electron detector (Gatan). In our setup, the dose protector was disabled for the K3 camera to be operated in diffraction mode. The camera was mounted on-axis at the bottom of the column, meaning the magnification by the projection lens system had to be adjusted for the longer physical distance between the sample and the detector. However, the projection of the beam stop, which is above the sample plane, takes up a large part of the detector and obscures many of the low-resolution reflections because of the longer physical distance. Therefore, in diffraction mode the beam stop was retracted for MicroED data collection. The sample to detector distance was a calibrated using a diffraction grating replica with amorphous gold shadowing (Ted Pella Inc., product no. 673). Compared to the K2 camera, the K3 has a larger field of view with 5,760×4,092 pixels. The camera operates internally at 1,500 Hz, with a maximum readout of 75 frames per second. The data rate of the K3 server in our setup, at reading out 40 fps (1× binning) in counting mode, is capped at 150 s, corresponding to an upper limit of 6,000 8-bit frames per movie. The linear range of the detector is approximately 15 e^-^/pixel/s at 90% DQE according to the manufacturer.

To ensure that all counts fall within the linear range of the camera we set up the microscope for low exposure diffraction data collection using the smallest C2 aperture (50 μm) and the largest spot size (11). The beam was spread to a diameter of 20 μm, corresponding to an exposure rate of approximately 0.0025 e^-^/Å^2^/s. MicroED data were collected at a set rotation speed of 0.15 °/s. A selected area (SA) aperture of 100 μm was used, corresponding to a diameter of 2 μm on the sample, to further improve the signal and minimize any background scattering from the surrounding area. For proteinase K lamellae, data were collected using a detector distance of 745 mm over a rotation range of 84° with a total exposure of 1.4 e^-^/Å^2^. Data were recorded using a total exposure of 420 s at either 2 or 10 fps. For triclinic lysozyme, data were collected using a detector distance of 373 mm over a rotation range of 63° with a total exposure of 1.05 e^-^/Å^2^. Data were recorded using a 560 s exposure at a frame rate of 10 fps. Electron-counting data were recorded using the K3 camera (1× binning, correct defects) and saved as 1-byte unsigned images in Gatan DM4 format using GMS (Gatan).

### 2.5 Data conversion and post-processing

Frames were converted from DM4 to MRC format within GMS. Gain normalization was applied during post-processing of the K3 data using the program clip in the IMOD software package (Kremer *et al*., 1996). The gain-normalized images were multiplied by a factor of 32 to facilitate data processing that requires integer valued pixels. Frames were converted from MRC to SMV format and summed using the MicroED tools (https://cryoem.ucla.edu). A total of 8 frames were summed for the K2 data collected at 40 fps. For the K3 data recorded at 10 fps, 5 frames were summed. No frames were summed for data collected with a readout of 2fps. The near-noiseless readout of an electron-counting detector implies that there is no penalty for dividing the rotation range into many small slivers instead of measuring reciprocal space in a few wide wedges. Individual frames can be summed into appropriately sized wedges after data collection. This is beneficial because it the optimal data collection strategy may not be known when a new sample is first inserted into the TEM.

### 2.6 Data processing and structure refinement

The diffraction data were indexed and integrated using XDS (Kabsch, 2010). Data were scaled using XSCALE (Kabsch, 2010), and merged in AIMLESS (Evans & Murshudov, 2013). The structures were determined by molecular replacement in Phaser using electron scattering factors (McCoy *et al*., 2007). Models were inspected and rebuild in Coot (Emsley *et al*., 2010), and the structures were refined with REFMAC5 using electron scattering factors (Murshudov *et al*., 2011). Correlation coefficients between Fo and Fc as function of resolution were calculated using the program sftools (Winn *et al*., 2011).

### 2.7 Figure and table preparation

Figures were made using matplotlib in Python 3.7 and ChimeraX. Figures were arranged in PowerPoint.

## 3. Results and Discussion

### 3.1 MicroED data collection using the K2

We explored the feasibility of collecting MicroED data in electron counting using a K2 direct electron detector. Data were collected from proteinase K crystals dispensed directly on the grid from the crystallization drop. Crystals were identified in search mode in overfocus diffraction phase contrast imaging manually brought to eucentric height. The projection of the beam stop on the K2 camera was obscuring much of the view and was therefore inserted only halfway (Fig. 1A). The white triangle in Fig 1A is the very tip of an enlarged beam stop. MicroED data were collected from three crystals that were rotated at a speed of 1.52°/s over wedges of 38.0°, 48.6°, and 60.8°. Data were recorded on the K2 camera in electron counting mode using 2× binning and 40 fps read out. Frames were hardware cropped with to a rectangle of 1650×1479 pixels improving the readout speed. The total exposure used for each dataset was less than 0.4 e^-^/Å^2^. As individual frames were sparse, frames were summed by 8 for each diffraction image during data conversion. Diffraction spots extended to beyond 2.5 Å resolution (Fig. 1A). Individual peak profiles show relatively sharp spots although overall the data are quite noisy and have high background counts (Fig. 1B). The individual crystal datasets were each integrated using XDS up to a resolution of 2.1 Å (Fig. 1C). Data were merged and truncated at 2.5 Å resolution at a mean I/σI of approximately 0.6 and CC_1/2_ of ∼0.3 indicating a significant correlation between two random half sets (Table 1, Fig. 1C) (Karplus & Diederichs, 2012).. The structure was determined by molecular replacement and refined using electron scattering factors and isotropic *B*-factors (Table 1). The map at moderate resolution, shows that individual side chains are resolved although some parts of the map are relatively noisy (Fig. 1D).

**Figure 1.**
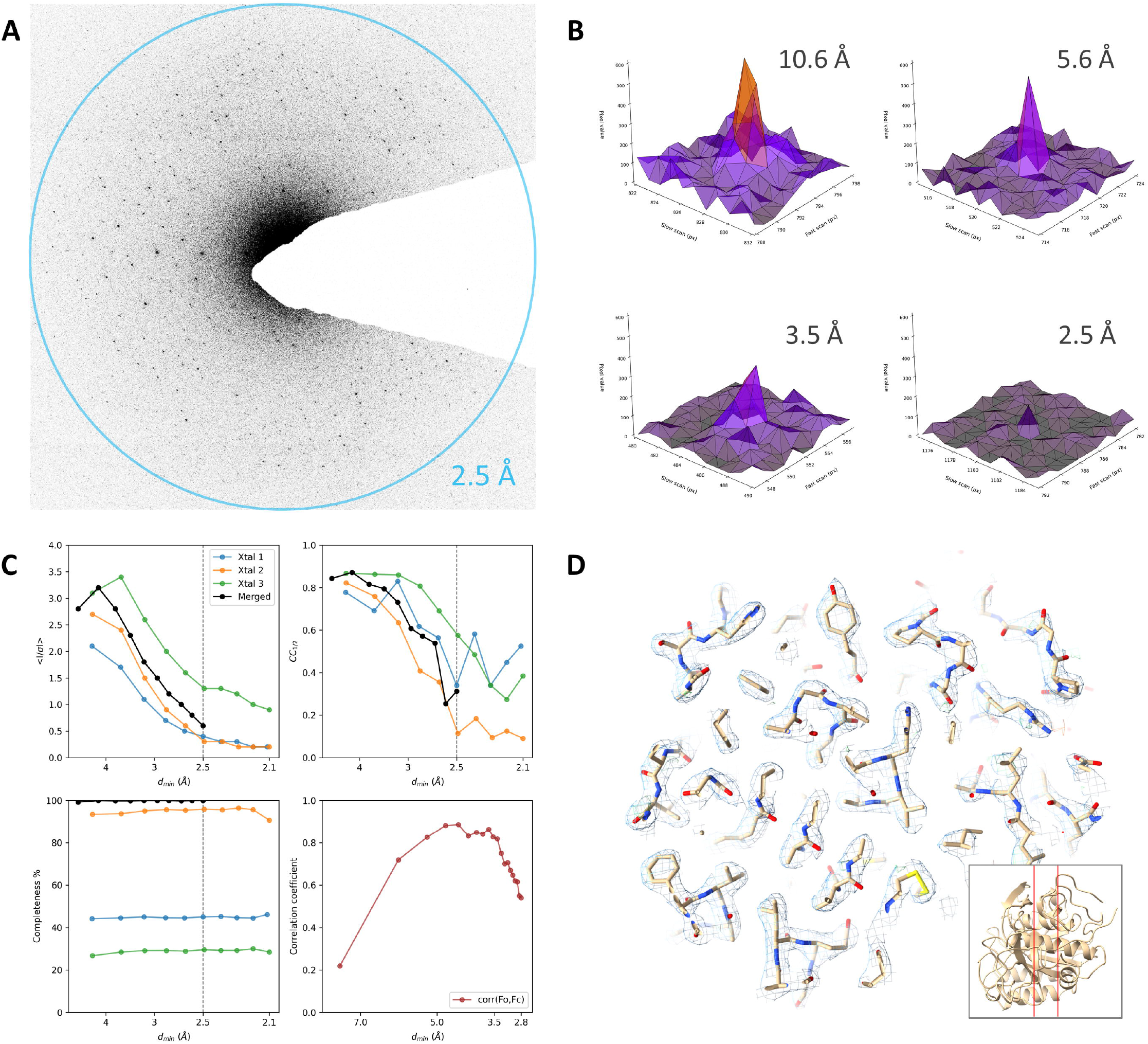
Electron-counting MicroED data of proteinase K using the K2 camera. (A) Diffraction pattern of a proteinase K crystal recorded using the K2 in electron counting mode, showing spots to beyond 2.5 Å resolution. For display, frames are cropped around the area of interest and were summed to cover a wedge in reciprocal space of approximately 1.0°. (B) Peak profiles at different resolutions are shown for individual frames used for data integration corresponding to a 0.3° wedge. (C) Plots showing the mean I/σI, CC_1/2_, and data completeness as function of the resolution for individual crystals datasets and the merged data. The fourth panel shows the correlation coefficient between the observed and calculated structure factor amplitudes for equal sized resolution bins. (D) The refined map shown for a slice through the structural model as indicated by the inset. For comparison, the same slice is slice shown in Fig. 2D. The 2mFo-DFc map is shown as blue mesh at a contour level of 1.2σ, the mFo-DFc difference map is contoured at ±3σ as green and red mesh for positive and negative values, respectively.

**Table 1.** Data processing and refinement statistics.

|  | Proteinase K (K2) | Proteinase K (K3) | Lysozyme (K3) |
| --- | --- | --- | --- |
| **Data collection** |  |  |  |
| Wavelength (Å) | 0.0197 | 0.0197 | 0.0197 |
| No. of crystals | 3 | 5 | 4 |
| **Data processing** |  |  |  |
| Space group | $P4_32_12$ | $P4_32_12$ | $P1$ |
| Unit cell dimensions |  |  |  |
| $a, b, c$ (Å) | 67.57, 67.57, 100.91 | 67.01, 67.01, 106.56 | 26.38, 30.76, 33.00 |
| $\alpha, \beta, \gamma$ (°) | 90, 90, 90 | 90, 90, 90 | 87.85, 108.85, 112.60 |
| Resolution (Å) | 25.92–2.50 (2.59–2.50) | 43.40–1.70 (1.73–1.70) | 31.08–1.20 (1.22–1.20) |
| Observed reflections | 66,710 (6,973) | 598,583 (11,074) | 98,413 (5,247) |
| Unique reflections | 8,816 (839) | 27,211 (1,289) | 24,986 (1,272) |
| Multiplicity | 7.4 (8.3) | 22.0 (8.6) | 3.9 (4.1) |
| Completeness (%) | 99.8 (100.0) | 99.3 (91.3) | 88.5 (90.1) |
| $R_{\text{merge}}$ | 0.807 (2.436) | 0.638 (1.574) | 0.215 (1.553) |
| $R_{\text{meas}}$ | 0.864 (2.607) | 0.653 (1.675) | 0.249 (1.780) |
| $R_{\text{pim}}$ | 0.290 (0.866) | 0.134 (0.551) | 0.122 (0.851) |
| Mean $I/\sigma I$ | 1.8 (0.6) | 4.8 (1.1) | 3.9 (1.0) |
| $CC_{1/2}$ | 0.814 (0.312) | 0.972 (0.108) | 0.984 (0.119) |
| **Refinement** |  |  |  |
| Resolution (Å) | 25.92–2.80 | 43.33–1.70 | 31.08–1.20 |
| No. reflections | 6,035 | 25,625 | 23,326 |
| No. reflections used for $R_{\text{free}}$ | 391 | 1,286 | 1,186 |
| $R_{\text{work}}/R_{\text{free}}$ | 0.240/0.296 | 0.176/0.254 | 0.181/0.242 |
| R.m.s. deviations |  |  |  |
| Bond lengths (Å) | 0.004 | 0.012 | 0.021 |
| Bond angles (°) | 1.354 | 1.430 | 2.142 |
| Mean $B$-factor (Å<sup>2</sup>) | 20.32 | 15.89 | 12.94 |
| Ramachandran |  |  |  |
| Favored (%) | 90.97 | 93.50 | 94.49 |
| Allowed (%) | 7.22 | 5.78 | 5.51 |
| Outliers (%) | 1.81 | 0.72 | 0.00 |
| Rotamer outliers (%) | 0.36 | 0.67 | 0.78 |

\*Values in parentheses are for highest-resolution shell. Data were truncated at a mean $I/\sigma I$ of approximately 1.0 and a still significant cross correlation between two random half sets at the 0.1% level (Karplus & Diederichs, 2012)

We demonstrate MicroED data collection using electron counting mode on the K2. Whereas a nearly complete dataset can be obtained and the structure can be solved at 2.5 Å resolution, the results are not optimal and several improvements are desirable. For example, owing to the position of the camera at the bottom on the column, the projection of the beam stop used in diffraction mode takes up quite a substantial part of the detector area, even when only inserted half-way (Fig. 1A). As a result, many reflections will be blocked by the beam stop and are not measured. This is not ideal, but not an insurmountable problem for diffraction, as symmetry equivalent reflections or Friedel mates would be observed opposite from the beam stop, but can be more challenging for lower symmetry space groups limiting data completeness. The frames recorded using the K2 do show different chip areas with seemingly different contrast. These stripe-like artifacts can likely be attributed to non-optimal normalization of the detector related to the framerate, and faster processing may lead to better application of the dark and gain reference tables. Background counts in the K2 images were still relatively high (Fig. 1A,B). Frames were recorded using hardware binning and were summed in post-processing, which would in part explain the background noise level. The data are of sufficient quality to solve and refine the structure although the resulting model *R*-factors are relatively poor (Table 1). Furthermore, the correlation between observed and calculated structure factor amplitude shows a poor correlation for both the lower and higher resolution reflections (Fig. 1C). On average, the low resolution reflections are more intense and the poor correlation indicates these counts may have been capped, even when using a low exposure rate and fast readout to minimize coincidence loss. Additional factors that can contribute to the noise at lower resolution also include the increased diffuse background close to the central beam owing to inelastic scattering (Fig. 1A). The higher resolution reflections are relatively weak and the poor correlation can be attributed to the background noise. Finally, the speed of the rotation and of the camera coupled with the large images and limits on available memory meant that only small wedges could be recorded on this particular system, even when using 2× binning and hardware cropping.

### 3.2 MicroED data collection using the K3

The K3 direct electron detector has an almost four times faster internal frame rate compared to the K2, meaning an improved linear response per pixel and expected less coincidence loss at similar exposure. To test the performance of the K3 in electron counting mode for MicroED experiments we collected data from crystal samples of proteinase K and triclinic lysozyme. On our system, the dose protector had to be disabled to allow the K3 camera to be inserted when the microscope is in diffraction mode. Furthermore, the beam stop was retracted in our setup as its projection was shading a large part of the detector area. The center beam is thus directly hitting the detector, enabling very accurate focusing using the live view of the camera. Damage to the camera caused by the direct beam, or highly intense low-resolution Bragg spots, is often cited as a major concern when using these types of detectors in diffraction. During our experiments, having low resolution spikes and even the center beam directly hitting the camera did not appear to cause any lasting damage to the detector. However, it may be that prolonged exposure to the direct beam could degrade the performance and reduce the lifetime of the central area of the camera. To avoid bias from burn-in, the camera was annealed, and new dark and gain references were taken prior to each set of experiments.

To optimize the signal-to-noise ratio for MicroED experiments the crystals were machined into thin lamellae using a dual-beam FIB/SEM. This improves the signal-to-noise ratio and also remove background scattering from the carbon support layer. Five crystals of proteinase K and an additional four crystals of triclinic lysozyme were identified for thinning using SEM. These crystals were thinned using a beam of gallium ions to the desired thickness and samples were cryo-transferred to the TEM directly after milling. Grids were aligned such that the milling direction is (close to) perpendicular to the rotation axis on the TEM. Lamellae sites were identified in the TEM from a whole-grid atlas. Each crystal lamella was manually brought to eucentric height and the positions were stored. Diffraction was focused and aligned on the K3 camera using live view. Before data collection, diffraction of each lamella was inspected by taking a single 2 s exposure in electron counting mode at a tilt angle of 0°.

Continuous rotation MicroED data of proteinase K were collected from wedges of -31.5° to +31.5° with a total exposure time of 420 s. The total exposure used was about 1.4 e^-^/Å^2^. Data were collected from 5 crystal lamellae and diffraction spots were observed up to 1.7 Å resolution (Fig. 2A). The Bragg peaks are very sharp and can be well distinguished above the background counts that are generally very low and far less noisy compared to earlier experiments using unmilled crystals with the K2 (Fig. 1A,B and Fig. 2A,B). The data were merged, reaching an overall completeness of approximately 99% at 1.7 Å resolution (Table 1, Fig. 2C). The higher resolution reached on the K3 can be attributed to the better signal-to-noise ratio from crystal lamellae where most surrounding ice and non diffracting material has been removed, and more accurate electron detection owing to the improved readout speed compared to the K2. The correlation plot for the K3 data shows a strong correlation for the low-resolution reflections, indicating that there are no apparent issues related to capping of the intensities (Fig. 2C). The resulting maps are of high quality and shows much more well-defined structural features compared to the proteinase K structure from the K2 data at poorer resolution (Table 1, Fig. 1D, Fig. 2D).

**Figure 2.**
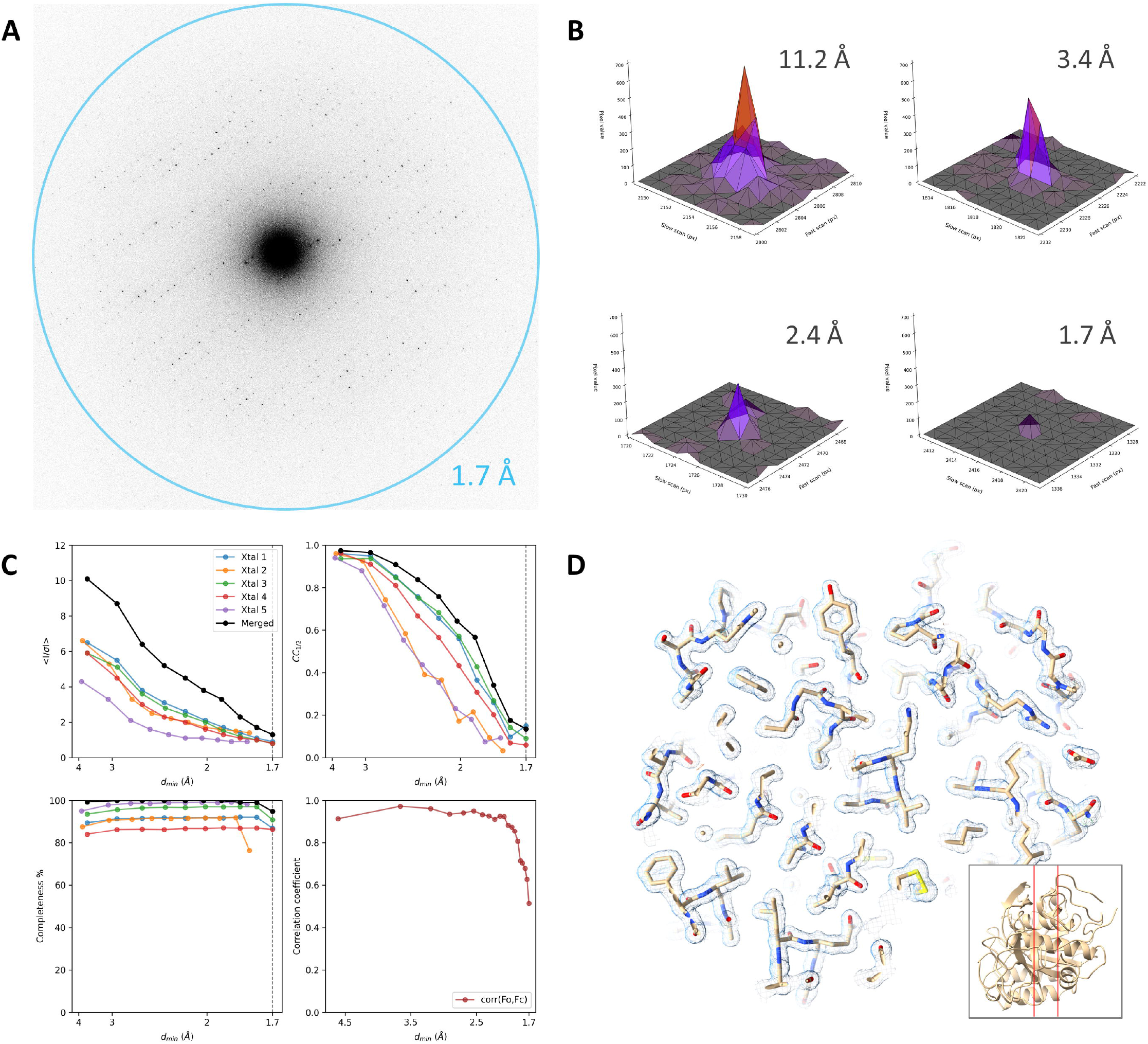
Electron-counting MicroED data of proteinase K using the K3 camera. (A) Diffraction pattern of a proteinase K lamella recorded using the K3 in electron counting mode, showing spots up to 1.7 Å resolution. For display, frames are cropped around the area of interest at the diffraction limit and were summed to cover a wedge in reciprocal space of approximately 1.0°. (B) Peak profiles at different resolutions are shown for individual frames used for data integration corresponding to a 0.076° wedge. (C) Plots showing the mean I/σI, CC_1/2_, and data completeness as function of the resolution for individual crystals datasets and the merged data. The fourth panel shows the correlation coefficient between the observed and calculated structure factor amplitudes for equal sized resolution bins. (D) The refined map shown for a slice through the structural model as indicated by the inset. For comparison, the same slice is shown in Fig. 1D. The 2mFo-DFc map is shown as blue mesh at a contour level of 1.2σ, the mFo-DFc difference map is contoured at ±3σ as green and red mesh for positive and negative values, respectively.

MicroED data of triclinic lysozyme were collected from a wedge of -42° to +42° with a total exposure time of 560 s. The total exposure used was approximately only 1.05 e^-^/Å^2^. Datasets were collected from 4 crystal lamellae and diffraction spots extended to 1.0 Å resolution (Fig. 3A). Similar to the proteinase K datasets, the background counts on the K3 camera are very low and the Bragg peaks are well defined (Fig. 3B). Whereas completeness of individual datasets is quite low owing to the low symmetry space group, data were merged reaching a completeness of 89% overall at 1.2 Å resolution (Fig. 3C). At high resolution, the structure is very well defined and individual atoms can be distinguished (Fig. 3D). Even though some of the lower resolution spots for the triclinic lysozyme data can be broad and very intense, there appears to be no sign of capping for these intensities judging from the correlation plot (Fig. 3C).

**Figure 3.**
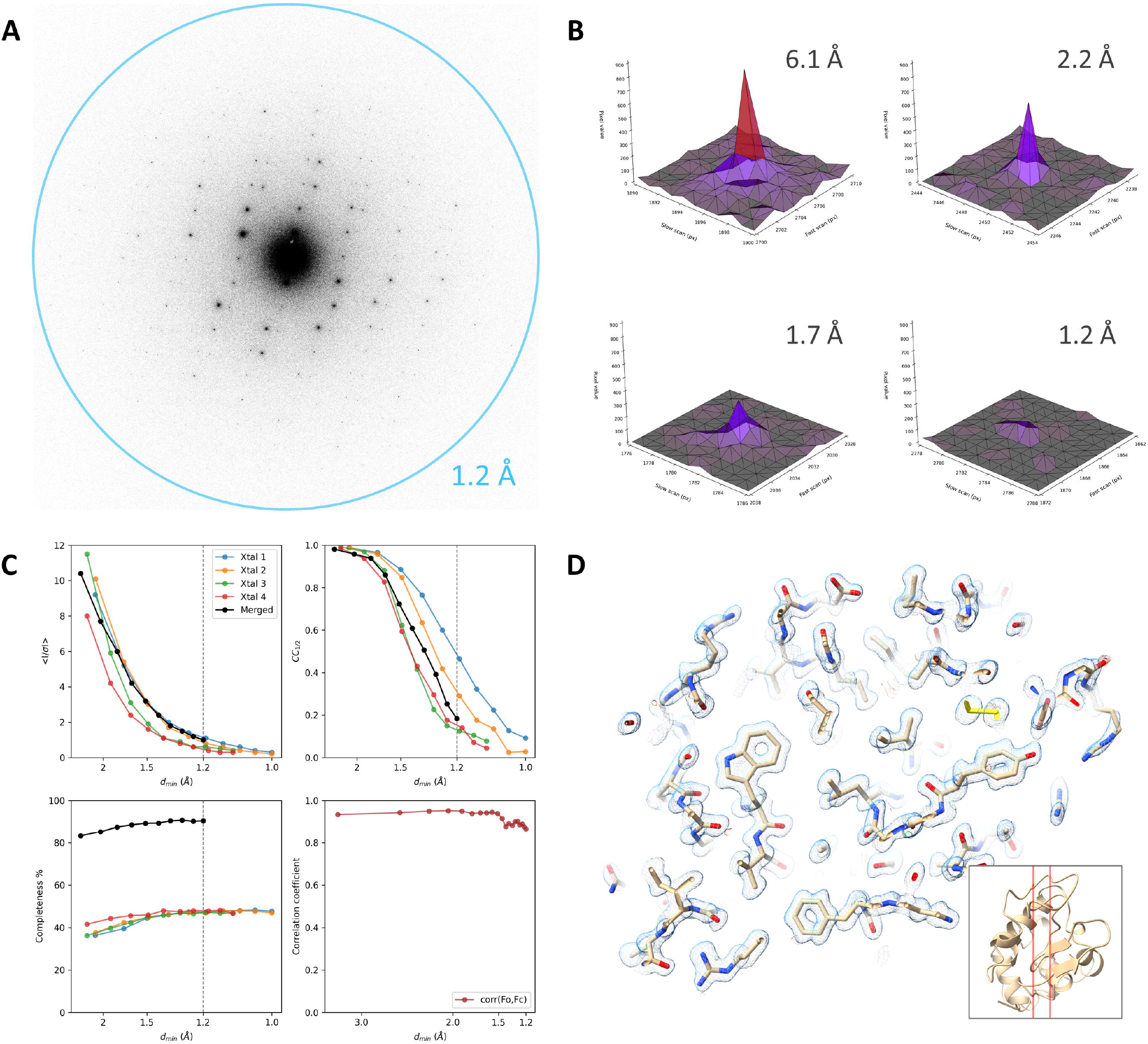
Electron-counting MicroED data of triclinic lysozyme using the K3 camera. (A) Diffraction pattern of a lysozyme lamella recorded using the K3 in electron counting mode, showing spots to beyond 1.2 Å resolution. For display, frames are cropped around the area of interest at the resolution limit and were summed to cover a wedge in reciprocal space of approximately 1.0°. (B) Peak profiles at different resolutions are shown for individual frames used for data integration corresponding to a 0.076° wedge. (C) Plots showing the mean I/σI, CC_1/2_, and data completeness as function of the resolution for individual crystals datasets and the merged data. The fourth panel shows the correlation coefficient between the observed and calculated structure factor amplitudes for equal sized resolution bins. (D) The map shown for a slice through the structural model as indicated by the inset. The 2mFo-DFc map is shown as blue mesh at a contour level of 1.2σ, the mFo-DFc difference map is contoured at ±3σ as green and red mesh for positive and negative values, respectively.

## 4. Conclusions

We present macromolecular structures of proteinase K and lysozyme from MicroED data recorded using the K2 and K3 direct electron detectors in counting mode. Although data from either cameras yields structural models, the faster frame rate on the K3 made a clear improvement in terms of data quality and attainable resolution. Data were collected using a low exposure rate for each movie, ensuring that intensities could be accurately measured by staying well within the linear response of the camera. For both samples, the total number of electrons used is significantly lower than what was used with previous data collection using scintillator based CMOS cameras or direct electron detectors in integrating/linear mode (Nannenga *et al*., 2014; Hattne *et al*., 2019). Because of the higher efficiency of the detector, the diffraction data still yields complete datasets of lysozyme and proteinase K even though the number of electrons per frame is greatly reduced. Under these conditions, we demonstrate that the same direct electron detectors that are used in other cryo-EM modalities can effectively also be used for MicroED experiments and macromolecular structure determination. The quality of the MicroED data may be further improved by increasing framerate and dynamic range, or even an event-based approach to electron counting. Additionally, the use of an energy filter removing the inelastically scattered electrons can reduce the diffuse background and further sharpen the peak profiles (Yonekura *et al*., 2002, 2019). These results can make MicroED more accessible to a wider audience in structural biology as electron counting detectors are typically available in cryo-EM user facilities and can provide accurate intensities for protein structure determination without the need for dedicated cameras.

## Data availability

Coordinates and structure factors have been deposited to the PDB.

## Acknowledgements

The authors would like to thank Ana Pakzad for helpful discussions. This study was supported by the National Institutes of Health P41GM136508 and R01GM124152. The Gonen laboratory is supported by funds from the Howard Hughes Medical Institute.

## References

Clabbers, M. T. B., van Genderen, E., Wan, W., Wiegers, E. L., Gruene, T. & Abrahams, J. P. (2017). Acta Crystallogr. Sect. Struct. Biol. 73, 738–748.

Clabbers, M. T. B., Martynowycz, M. W., Hattne, J. & Gonen, T. (2022). Hydrogens and hydrogen-bond networks in macromolecular MicroED data. bioRxiv https://doi.org/10.1101/2022.04.08.487606

Cowley, J. M. & Moodie, A. F. (1957). Acta Crystallogr. 10, 609–619.

Emsley, P., Lohkamp, B., Scott, W. G. & Cowtan, K. (2010). Acta Crystallogr. D Biol. Crystallogr. 66, 486–501.

Evans, P. R. & Murshudov, G. N. (2013). Acta Crystallogr. D Biol. Crystallogr. 69, 1204–1214.

Hattne, J., Martynowycz, M. W., Penczek, P. A. & Gonen, T. (2019). IUCrJ. 6, 921–926.

Hattne, J., Shi, D., Glynn, C., Zee, C.-T., Gallagher-Jones, M., Martynowycz, M. W., Rodriguez, J. A. & Gonen, T. (2018). Structure. 26, 759-766.e4.

Henderson, R. (1995). Q. Rev. Biophys. 28, 171–193.

Kabsch, W. (2010). Acta Crystallogr. D Biol. Crystallogr. 66, 125–132.

Karplus, P. A. & Diederichs, K. (2012). Science. 336, 1030–1033.

Kremer, J. R., Mastronarde, D. N. & McIntosh, J. R. (1996). J. Struct. Biol. 116, 71–76.

Li, X., Mooney, P., Zheng, S., Booth, C. R., Braunfeld, M. B., Gubbens, S., Agard, D. A. & Cheng, Y. (2013). Nat. Methods. 10, 584–590.

Li, X., Zheng, S. Q., Egami, K., Agard, D. A. & Cheng, Y. (2013). J. Struct. Biol. 184, 251–260.

Martynowycz, M. W., Clabbers, M. T. B., Hattne, J. & Gonen, T. (2022). Nat. Methods. 19, 724–729.

Martynowycz, M. W., Clabbers, M. T. B., Unge, J., Hattne, J. & Gonen, T. (2021). Proc. Natl. Acad. Sci. 118, e2108884118.

McCoy, A. J., Grosse-Kunstleve, R. W., Adams, P. D., Winn, M. D., Storoni, L. C. & Read, R. J. (2007). J. Appl. Crystallogr. 40, 658–674.

Murshudov, G. N., Skubák, P., Lebedev, A. A., Pannu, N. S., Steiner, R. A., Nicholls, R. A., Winn, M. D., Long, F. & Vagin, A. A. (2011). Acta Crystallogr. D Biol. Crystallogr. 67, 355–367.

Nakane, T., Kotecha, A., Sente, A., McMullan, G., Masiulis, S., Brown, P. M. G. E., Grigoras, I. T., Malinauskaite, L., Malinauskas, T., Miehling, J., Uchański, T., Yu, L., Karia, D., Pechnikova, E. V., de Jong, E., Keizer, J., Bischoff, M., McCormack, J., Tiemeijer, P., Hardwick, S. W., Chirgadze, D. Y., Murshudov, G., Aricescu, A. R. & Scheres, S. H. W. (2020). Nature. 587, 152–156.

Nannenga, B. L., Shi, D., Leslie, A. G. W. & Gonen, T. (2014). Nat. Methods. 11, 927–930.

Nederlof, I., van Genderen, E., Li, Y.-W. & Abrahams, J. P. (2013). Acta Crystallogr. D Biol. Crystallogr. 69, 1223–1230.

Takaba, K., Maki-Yonekura, S., Inoue, S., Hasegawa, T. & Yonekura, K. (2021). Front. Mol. Biosci. 7, 612226.

Winn, M. D., Ballard, C. C., Cowtan, K. D., Dodson, E. J., Emsley, P., Evans, P. R., Keegan, R. M., Krissinel, E. B., Leslie, A. G. W., McCoy, A., McNicholas, S. J., Murshudov, G. N., Pannu, N. S., Potterton, E. A., Powell, H. R., Read, R. J., Vagin, A. & Wilson, K. S. (2011). Acta Crystallogr. D Biol. Crystallogr. 67, 235–242.

Yonekura, K., Ishikawa, T. & Maki-Yonekura, S. (2019). J. Struct. Biol. 206, 243–253.

Yonekura, K., Kato, K., Ogasawara, M., Tomita, M. & Toyoshima, C. (2015). Proc. Natl. Acad. Sci. 112, 3368–3373.

Yonekura, K., Maki-Yonekura, S. & Namba, K. (2002). Biophys. J. 82, 2784–2797.

Zhou, H., Luo, Z. & Li, X. (2019). J. Struct. Biol. 205, 59–64.

